# Genome-wide selection inference at short tandem repeats

**DOI:** 10.1101/2022.05.12.491726

**Authors:** Bonnie Huang, Arun Durvasula, Nima Mousavi, Helyaneh Ziaei-Jam, Mikhail Maksimov, Kirk E. Lohmueller, Melissa Gymrek

## Abstract

Short tandem repeats (STRs) comprising repeated sequences of 1-6 bp are one of the largest sources of genetic variation in humans. STRs are known to contribute to a variety of disorders, including Mendelian diseases, complex traits, and cancer. Based on their functional importance, mutations at some STRs are likely to introduce negative effects on reproductive fitness over evolutionary time. We previously developed SISTR (Selection Inference at STRs), a population genetics framework to measure negative selection against individual STR alleles. Here, we extend SISTR to enable joint estimation of the distribution of selection coefficients across a set of STRs. This method (SISTR2) allows for more accurate analysis of a broader range of STRs, including loci with low mutation rates. We apply SISTR2 to explore the range of feasible mutation parameters and demonstrate substantial variation in mutation and selection parameters across different classes of STRs. Finally, we show that *de novo* STR mutations tend to confer a greater selective burden compared to standing STR variation in the population and measure the relative burden of STRs vs. single nucleotide variants in a typical genome. Overall, we anticipate that the evolutionary insights gained from this study will be important for future studies of variation at STRs and their role in evolution and disease.

## Introduction

Short tandem repeats (STRs) are DNA sequences consisting of repeated 1-6 base pair motifs that comprise approximately 1.6 million loci in the human genome^1^. Due to their high prevalence in the genome and rapid mutation rates, variation in copy number at STRs represents a large portion of human genetic variation. Recent evidence supports a role for STRs in diverse biological processes that control gene regulation^2,3^ and contribute to a wide range of human traits^4^. Based on their functional importance, mutations at some STRs are likely to introduce detrimental effects on reproductive fitness. Understanding these fitness effects can provide insights into the role of STRs in evolution.

Previous studies have used multiple approaches to measure the effects of natural selection on STRs. Haasl and Payseur developed a detailed model of STR evolution including mutation, genetic drift, and natural selection at an STR implicated in Friedreich’s Ataxia^5^. However, fitting their model is computationally intensive due to the large number of parameters, making it infeasible to fit individually at each of the more than one million STRs in the genome. We previously developed an STR constraint metric based on comparing observed vs. expected mutation rates^6^. This metric could broadly distinguish neutrally evolving STRs from those implicated in severe early-onset disorders. However, that score is based on noisy STR mutation rates that are computationally expensive to estimate from individual-level genotypes, does not model the known dependence of mutation rate on allele length, and only produces locus-level, rather than allele-level scores.

Recently, we introduced SISTR (Selection Inference at Short Tandem Repeats), a computationally efficient method to measure negative selection at STRs^7^. SISTR estimates per-locus selection coefficients by finding the selection parameters that best fit the allele frequency distribution for one STR at a time. These fine-grained scores enable predicting the fitness impact of individual alleles at a specific locus. However, this approach faces several limitations. First, it has low power to detect weak selective effects. Furthermore, at STRs with extremely low mutation rates, such as short trinucleotide repeats, it is unable to precisely estimate selection coefficients since low levels of genetic variation could be due to the low mutation rate, strong negative selection, or a combination of both forces.

Here, we extend SISTR to enable joint estimation of the distribution of selection coefficients across a set of STRs. This method allows us to more accurately analyze a broader range of STRs, including loci with low mutation rates. We apply SISTR2 to explore the range of feasible mutation parameters and demonstrate substantial variation in mutation and selection parameters across different classes of STRs. Finally, we show that *de novo* STR mutations tend to confer stronger selective burden compared to standing STR variation in the population and measure the relative burden of STRs vs. single nucleotide variant (SNV) mutations in a typical genome. Overall, we anticipate that the evolutionary insights gained from this study, including a more detailed understanding of STR mutation and selection parameters for different types of STRs, will be important for future studies of variation at STRs and their role in evolution and disease.

## Results

### Overview and validation of the SISTR2 joint inference method

We previously developed SISTR, a population genetics framework that estimates selection coefficients at individual STRs^7^. SISTR incorporates an evolutionary model of STR variation that includes mutation, negative natural selection, and genetic drift. Our mutation model is based on a generalized stepwise mutation (GSM) model with two modifications, including a length-dependent mutation rate and a directional bias in mutation sizes toward an optimal (central) allele length (**Fig. 1a**). Since the optimal allele is typically unknown, for most applications we assume the modal allele in the population is the optimal allele and treat these interchangeably. To model negative selection, we assume that the modal allele at each STR has a fitness of 1, and that the fitness of other alleles decreases linearly with their distance, in number of repeat units, from the optimal allele. The decrease in fitness of non-optimal alleles scales with *s*, which ranges from 0 (no effect on fitness) to 1 (any allele other than the optimal allele is lethal). SISTR leverages a previously developed technique^5^ that incorporates mutation, selection, and demographic models (**Methods**) to simulate allele frequencies forward in time (**Supplementary Fig. 1**). Using approximate Bayesian computation (ABC), we determine the posterior distribution of *s* at each locus by comparing observed allele frequencies to those simulated by our model. We previously showed that SISTR performs well on STRs with high mutation rates, but is underpowered to detect selection at STRs with low mutation rates or under only modest selection. In those cases, information contained in genetic variation at a single locus is insufficient to accurately infer selection.

**Figure 1:**
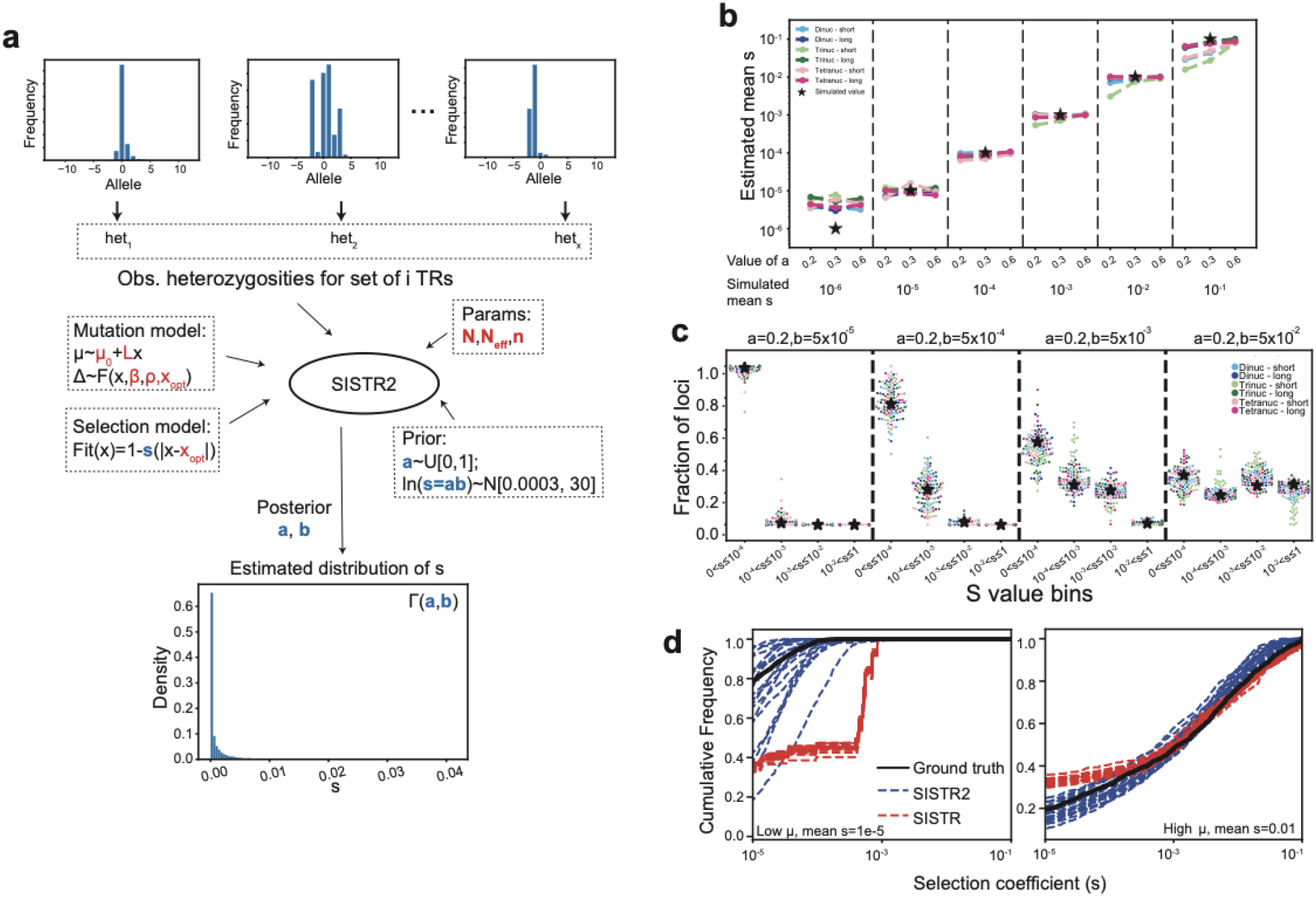
SISTR2 estimates the distribution of fitness effects (DFE) across a set of STRs. **(a) Overview of SISTR2**. For a set of STRs, SISTR2 takes priors on the parameters of the gamma distribution of fitness effects (*s*), a mutation model, a selection model, demographic parameters, and the observed distribution of heterozygosities across a set of STRs as input. It outputs a posterior estimate of the parameters *a* and *b* that describe the gamma distribution of fitness effects (DFE) across loci. Bolded red variables indicate input parameters describing the mutation, selection, and demographic models. The full model is described in **Methods**. **(b) Validation of SISTR2 using simulated data**. The x-axis indicates the simulated gamma distribution parameters. Dashed vertical lines separate simulation settings with the same mean *s* value. For a given mean *s* value, various simulations with different gamma distribution shapes (controlled by *a*) were run. The y-axis gives the estimated mean *s* of 20 estimates. **(c) Inferred distribution of *s* values for four gamma distribution parameter (*a, b*) combinations**. The x-axis denotes bins of *s* values. The y-axis gives the fraction of loci inferred to be in each bin. The black stars give the ground truth fraction of *s* values in each bin. For **(b)** and **(c)** each color represents a different mutation model setting (**Methods**). **(d) Comparison of DFEs inferred by SISTR (per-locus) vs. SISTR2 (joint)**. Plots show the cumulative frequency distribution of *s* for various simulation rounds estimated either individually at each locus using SISTR (red) or inferred jointly across all loci using SISTR2 (blue). The black line shows the ground-truth distribution of *s* values. The left panel shows a setting with low mutation rate and low selection (mean *s*=1e-5), highlighting a case where SISTR, but not SISTR2, overestimates *s*. The right panel shows a setting with high mutation rate and high selection (mean *s*=0.01), where both methods perform well. In both settings, SISTR2 produces unbiased estimates of the distribution of *s* values.

To address these challenges, we developed SISTR2, an extension of SISTR that enables joint estimation of the distribution of fitness effects (DFE) across a set of STRs (**Methods; Fig. 1a; Supplementary Fig. 2**). By leveraging information across a set of loci, SISTR2 can obtain more precise estimates of the DFE. Instead of estimating *s* at each STR individually, the joint method assumes *s* for each STR is drawn from a gamma distribution parameters of this distribution. *s* ~ Γ(*a, b*) and infers the parameters of this distribution.

SISTR2 takes as input allele frequencies for a set of STR loci, prior distributions on the gamma distribution parameters, and the mutation, selection, and demographic models used as input to SISTR. It first computes heterozygosity (defined as 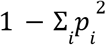, where *p*_*i*_ is the frequency of the *i* th allele) as a summary statistic for each locus (**Methods**). It then uses our simulation framework to simulate a set of STR loci such that the selection coefficient for each locus is drawn from Γ(*a, b*) for some *a* and *b*. Using ABC, SISTR2 determines the median posterior estimates of *a* and *b* by comparing heterozygosity distributions of observed vs. simulated data for different values of *a* and *b*.

Using simulated data, we validated the ability of the joint method implemented in SISTR2 to obtain a posterior estimate of the gamma distribution parameters describing a DFE for a set of STRs. We tested the method on different mutation models as well as a variety of gamma distribution parameters capturing a range of distributions of selective effects. In each simulation, we first chose a mutation model and optimal allele length, then simulated STR allele frequencies for 1,000 loci, drawing the value of *s* for each locus from a gamma distribution. We then used SISTR2 to estimate *a* and *b* from each simulated dataset and compared inferred values to the true values used to simulate the data (**Methods**). We found SISTR2 accurately recovered simulated gamma distribution parameters for mean *s* values ranging from 10^−5^ to 10^−1^, that our point estimates of *s* are unbiased (**Fig. 1b**), and that inferred distributions of *s* match well to the true distributions used to simulate the data (**Fig. 1c**). Further, these estimates are robust to common STR genotyping errors (**Supplementary Fig. 3, Methods**).

Next, we compared joint estimates of *s* obtained from SISTR2 to per-locus estimates output by SISTR. To this end, we used SISTR to estimate individual *s* values for each simulated locus above, and compared the distribution of *s* values to those from the true underlying gamma distribution (**Fig. 1d, Supplementary Fig. 4**). We found that joint estimation with SISTR2 shows several important advantages. First, it can distinguish between weaker selection (e.g. *s* = 10^−5^ vs. 10^−4^) and neutrality (*s*=0) which is difficult to do using SISTR. Second, it can more accurately infer selection at loci with low mutation rates. We further evaluated the impact of the number of loci used on SISTR2’s performance. As expected, precision increases with the number of loci used as input to the joint estimation. For most settings, accurate estimates of *s* can be obtained with as few as 10 loci (**Supplementary Fig. 5**). For sets of loci with lower mutation rates (<10^−5^), several hundred loci are needed.

### Inferring feasible mutation parameters for various types of STRs using SISTR2

Accurate models of the mutational process are necessary to infer selection on putatively functional STRs. In addition to inferring DFEs, SISTR2’s framework allows us to determine the range of feasible mutation parameters for a set of neutrally evolving loci. To do so, we draw mutation rates from a uniform prior, fix *s*=0, and run SISTR2’s ABC method on a target set of presumed neutral loci. If the posterior distribution of accepted mutation rates is wide, it indicates that a broad range of mutation models are consistent with the observed heterozygosity distribution or that there is not enough data available to determine which mutation model feasibly explains the observed data. On the other hand, if the posterior distribution is tight, it indicates that only one mutation model fits best.

We performed STR genotyping using GangSTR^8^ in 534 samples of European descent (**Discussion**) from the 1000 Genomes Project^9^ for which deep whole-genome sequencing data was available^10^ (**Methods**). We restricted our analysis to STRs with repeat unit lengths 2-4 bp, which are abundant in the genome and can be reliably genotyped. We further filtered very short STRs (**Methods**) since those loci are typically not polymorphic in repeat copy number. After filtering, 86,327 STRs remained for analysis. We then applied SISTR2 using the strategy described above to explore feasible ranges for mutation parameters across STRs with a range of repeat unit lengths, sequences, and modal (optimal) allele lengths. For dinucleotides, we tested 6 different mutation rate models (**Fig. 2a**), and for trinucleotides and tetranucleotides, we tested 7 models (**Fig. 2b and 2c**), each of which models a linear relationship between repeat length and mutation rate. For each setting, we applied SISTR2 to intergenic loci which we expect are mostly neutral, and set the gamma distribution parameters *a* and *b* such that the *s* value drawn is always 0. Then, we recorded which mutation models were accepted via ABC for each class of loci and assessed whether accepted simulations were enriched for particular mutation models (**Fig. 2d-f**). The model with the highest number of ABC acceptances corresponds to the maximum likelihood mutation model.

**Figure 2:**
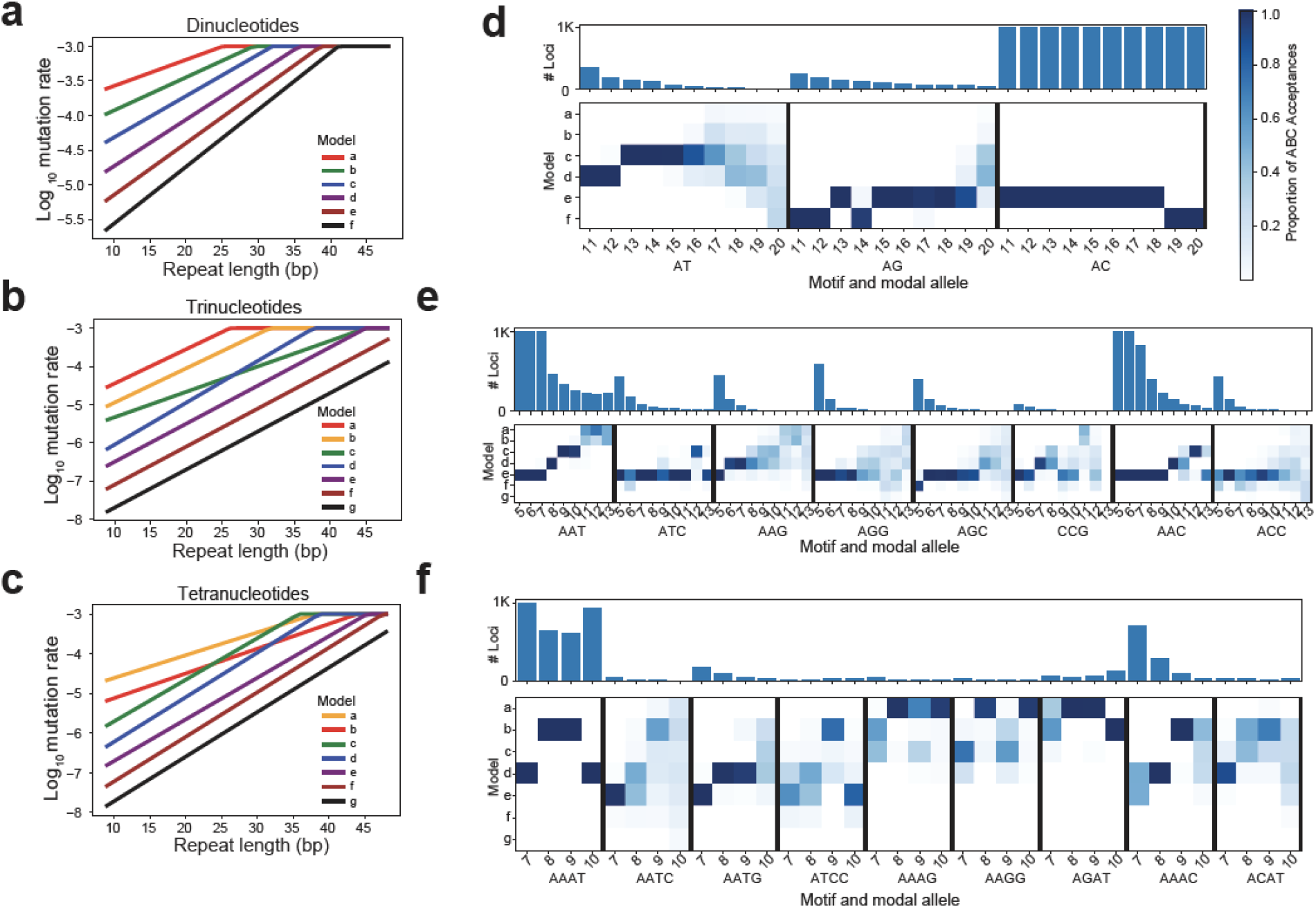
Exploring feasible mutation rates consistent with observed allele frequencies. **(a-c) Mutation rate models tested for dinucleotide, trinucleotide, and tetranucleotide STRs**. The x-axis shows the allele length in base pairs, and the y-axis gives the corresponding log_10_ mutation rate. **(d-f) Mutation model fit for different STR classes**. Each STR class (x-axis) is defined by the repeat sequence and modal (optimal) allele length. Top panels denote the number of loci in each class (truncating at 1,000 for categories with more loci). For bottom panels, each row denotes one mutation setting corresponding to models shown in **a-c**. The color of each cell represents the proportion of the ABC acceptances each mutation model represents.

We assessed the goodness of fit of each model using a post-hoc Kolmogorov-Smirnov (KS) test to compare distributions of heterozygosity values for loci simulated under the model to the observed heterozygosity distribution for a set of STRs (**Supplementary Fig. 6**). We repeated this test 100 times for each class of STRs and recorded the percentage of simulation rounds where the observed distributions were significantly different (nominal KS p<0.05). Notably, STR classes with the highest numbers of loci have the highest power to detect even small differences in heterozygosity distributions. To reduce this bias in power, we repeated the analysis after subsetting the number of loci in each class to a maximum of 50 STRs (**Supplementary Fig. 7**). For most classes of loci, the percentage of simulations that are similar to the observed values is high, indicating the maximum likelihood mutation model fits well. However, for certain classes of loci (e.g. long AC repeats, short trinucleotide repeats), simulated and observed heterozygosity distributions were significantly different (KS p<0.05). We plotted examples of observed versus simulated heterozygosity distributions to visually assess their similarity (**Supplementary Fig. 8**). We found in most cases, the observed heterozygosity distribution is still largely similar to that produced by the inferred maximum likelihood model. Overall, these results suggest that there may be aspects of the mutational process at these loci that are not captured by our model, but that the maximum likelihood mutation model still provides a reasonable fit for use in downstream analyses.

We found that even among loci with the same repeat unit length, there is notable variation in feasible mutation rates across different repeat unit sequences. For example, within dinucleotides, we found that a single mutation model fits most loci with repeat units AC and AG well, whereas our analysis suggests most AT repeats have higher mutation rates (**Fig. 2d**). This trend is consistent with mutation rates we^7^ and others^11^ previously inferred from *de novo* STR mutations, which found that STRs with repeat unit AT mutate several times faster than other dinucleotides STRs (**Supplementary Fig. 9**).

Most trinucleotide repeats are fit by a single mutation model, with some notable exceptions (**Fig. 2e**). For example, AAT repeats with modal alleles greater than 8 repeat units fit best with higher mutation rates than expected even accounting for a linear increase of mutation rate with repeat length. One explanation for this deviation could be that mutation processes at long AAT repeats are not captured by our model and that mutation rates scale super-linearly with repeat length. An alternative explanation is that intergenic AAT repeats may not be truly neutrally evolving, which could bias our mutation model inference. Within trinucleotides, we also estimate that AAG have consistently higher mutation rates, which matches previous observations that these repeats are unstable and prone to large expansions^12^.

Maximum likelihood mutation models for tetranucleotides showed more variability across repeat unit sequences, and are consistent with previous reports that AAAG, AAGG, and AGAT repeats exhibit higher mutation rates than other tetranucleotide STRs^12^. Overall, this analysis highlights substantial differences in mutation rate across STRs with different repeat units. We used these results to inform repeat unit-specific mutation rate parameters for selection inference performed below.

### Distinct types of repeats have different fitness effects

Next, we applied SISTR2 genome-wide to estimate the DFE at different STR classes. As inputs, we used allele frequencies from 86,327 STRs computed from European samples as described above, and set mutation parameters at each STR based on the maximum likelihood mutation model for each repeat unit sequence and optimal allele identified in **Fig. 2**. We first examined DFEs for different repeat classes and functional categories weighted by the number of loci for each optimal allele (**Fig. 3a, Methods**). We found, as expected, that coding STRs are under strongest selection whereas intergenic STRs are under the weakest selection (**Fig. 3a**). Intronic STRs and those in UTRs or promoters are under slightly stronger selection than STRs in intergenic regions. Full results for each category are summarized in **Supplementary Table 1**.

**Figure 3:**
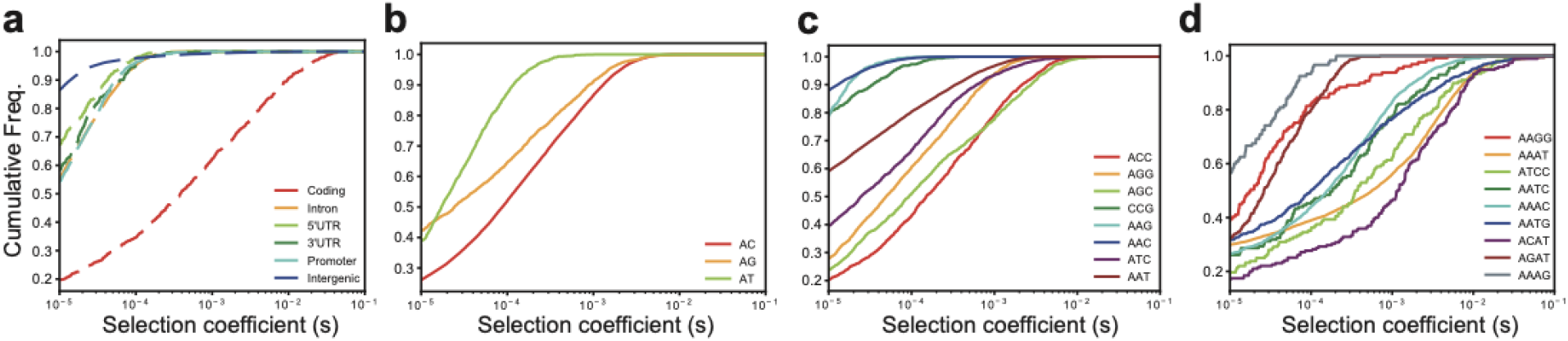
The distribution of selection coefficients for different STR classes. **(a) Functional category analysis**. Red=coding, orange=introns, light green=5’UTR, dark green=3’UTR, cyan=promoters; purple=intergenic. **(b-d) Repeat unit analysis**. For all plots, the CDF is weighted by the number of loci for each modal allele (**Methods**). Each color denotes a different repeat unit. All analyses presented here use the joint inference method implemented in SISTR2.

Next, we compared DFEs for STRs with different repeat unit sequences and optimal allele lengths (**Fig. 3b-d; Supplementary Table 2**). As in our mutation rate analysis, we tested the goodness of fit of the DFE estimated by SISTR2 using a KS-test and found that in most cases inferred DFEs resulted in good fits to observed heterozygosity distributions (**Supplementary Fig. 10**). Similar to our mutation rate analysis, observed data at most repeat classes fit well to SISTR2’s inferred model with several exceptions. In particular, short trinucleotides, AT repeats, and AAT repeats were not fit well, indicating inferred DFEs for those loci may be unreliable (**Discussion**).

Overall for dinucleotides, AC repeats tend to be under the strongest selection **(Fig. 3b)**. However, this trend is dependent on the length of the STR. Analyzing STRs separately by optimal allele length shows that longer AC repeats tend to be under increased selection, whereas for shorter STRs AG repeats are under strongest selection (**Supplementary Fig. 11**). We additionally observed that AT repeats tend to be under weaker selection than AC or AG repeats, although this trend may be driven by the overall shorter average lengths of AT vs. AC repeats (**Fig. 2**) or could reflect the relatively poor fit of our model to AT vs. other dinucleotide repeats (**Supplementary Fig. 10**). For trinucleotides, we found that ACC and AAC repeats tended to be under the strongest and weakest selection, respectively (**Fig. 3c**; **Supplementary Fig. 11**). For tetranucleotides, AAGG, AAAG, and AGAT, which also showed higher mutation rates than other tetranucleotide STRs^12^ (**Fig. 2f**), exhibited the weakest selection coefficients (**Fig. 3d**; **Supplementary Fig. 11**).

### Overall burden of deleterious mutations due to STRs vs SNVs

We used SISTR2 to quantify the genome-wide fitness burden of *de novo* STR mutations using two different methods (predicted and directly observed) (**Table 1**). In each case, we assumed the total mutation burden is additive across individual mutations and excluded STR mutation classes which had poor model fit based on the KS-test described above (**Supplementary Fig. 12, Methods**). This excludes a large number of AAAT and long AC repeats, which likely have deleterious effects but had unreliable estimates for *s*. First, we computed the predicted burden of mutations within each STR class separately based on their mutation rates, inferred DFEs, and mutation properties, and summed the burden across all classes. Second, we computed the burden of directly observed *de novo* STR mutations in 3 trio families from the 1000 Genomes Project with available deep WGS (**Methods**) by summing the predicted fitness effect of each mutation relative to the parent allele as computed by SISTR2. Notably, since WGS is based on cell-line derived DNA, some observed mutations are likely to be cell line artifacts rather than true *de novo* mutations. Both methods for computing the fitness burden of *de novo* STR mutations gave similar results (mean additive burden ranging from 0.01-0.03).

**Table 1:** Comparison of estimated fitness burden from de novo STR vs. SNV mutations. Burden refers to the sum of fitness effects across all mutations in each category. SNP burdens are computed based on average mutation rates and fitness effects for each class (**Methods**). For STRs, the expected burden is summed across the burden from each STR class (repeat unit and optimal allele). 95% confidence intervals were computed based on 100 bootstrap samples of mutations sampled from each class based on their relative proportions and sampling their selection coefficients from the gamma distribution inferred by SISTR2 (**Methods**). For observed mutations, 95% confidence intervals were computed based on 100 bootstrap samples of observed mutations. ^+^STR counts are considering only repeats with units 2-4bp that have repeat units common enough to be estimated by SISTR2 and with relatively good model fits (**Supplementary Table 2**). ^*^For observed de novo mutations, only mutations predicted to have negative fitness consequences, and for which SISTR2 scores were computed, were considered. Fitness effects for *de novo* mutations were computed as the difference in fitness between the new allele and the parent allele.

| Mutation type | Mean $s$ | # Mutations/generation | Total burden |
| --- | --- | --- | --- |
| STRs - genome-wide<sup>+</sup> |  |  |  |
| Predicted | 0.00077 | 13.78 | 0.012 (0.00080-0.039) |
| Observed (3 trios<sup>+</sup>) |  |  |  |
| NA12864 | 0.00051 | 41 | 0.021 (0.011-0.033) |
| NA10865 | 0.00055 | 58 | 0.032 (0.017-0.045) |
| NA10845 | 0.00088 | 17 | 0.015 (0.0048-0.028) |
| SNVs - missense | 0.0066 | 0.56 | 0.0037 |
| SNVs - conserved non-coding | 0.00056 | 2.54 | 0.0014 |

We next compared the burden of genome-wide *de novo* STR vs. SNV mutations (**Table 1**). For SNVs, we considered two classes of mutations for which mutation rates and fitness effects were available^13,14^: nonsynonymous mutations and mutations occurring in conserved non-coding regions (**Methods, Supplementary Table 4**). The expected fitness effect of an individual single-nucleotide missense mutation (*s*=6.6e-3) is approximately 10 times higher than that of an individual STR mutation (*s*=7.7e-4), whereas the effect of an individual single-nucleotide mutation in conserved non-coding regions (*s*=5.6e-4) is similar to that of an STR mutation. However, each individual genome is expected to have far more *de novo* STR mutations (mean=14, considering genome-wide mutations at STRs with repeat units 2-4bp and reliable selection models) compared to mutations resulting in missense SNVs (mean=0.56) or SNVs in conserved non-coding regions (mean=2.5). Thus overall, the expected total burden of genome-wide STR mutations is notably higher than either category of single-nucleotide mutations (approximately 3.2x and 8.6x higher than missense and conserved non-coding, respectively) and is higher than both categories of single-nucleotide mutations together.

The results above may overestimate the relative burden of STR vs. SNV mutations since it considers genome-wide STRs but only a subset of possible SNV mutations collectively covering less than 5% of the genome. We considered the theoretical mutation burden based on varying the proportion of non-coding single-nucleotide mutations under selection from 5% to 30% (**Supplementary Table 4**), and found that the overall burden in the most extreme case (8.57E-03 for 30% of the non-coding genome under selection) is still substantially smaller than the genome-wide STR burden (0.012). Overall, while this analysis faces several limitations (**Discussion**), our results suggest that although STR mutations individually have modest fitness effects, the larger number of STR mutations per generation results in a total fitness burden that is several times higher than that of missense, and similar or higher in magnitude to the genome-wide burden of all single-nucleotide mutations.

Finally, we compared the relative burden of inherited STRs vs. SNVs in a typical genome (**Supplementary Tables 5-6**). For STRs, we computed the fitness effects of inherited variation in the children of each of the three trios analyzed above summed across all non-optimal STR alleles (**Methods**). Similar to *de novo* mutations, we excluded repeat classes with poor model fit. As expected, inherited STR variants tended to have lower selection coefficients compared to *de novo* variants (**Supplementary Fig. 13**; KS two-sided p<0.01 for all samples). For SNVs, we considered observed variants in two classes: nonsynonymous variants and variants falling in conserved non-coding regions as measured by CADD^15^ score >15, corresponding to the top 3% of genome-wide scores (**Methods**). The estimated STR burden is substantially lower than that for SNVs in these categories (mean STR burden 8.81 excluding long AC/AAAT repeats, and mean SNV burden 27). The majority of this SNV burden is attributed to nonsynonymous SNVs (20 vs. 6.8 for conserved non-coding). Similar to our estimates for *de novo* mutations, we additionally varied the proportion of non-coding variants potentially under selection by considering a range of CADD score thresholds. For more permissive definitions of conserved non-coding regions (CADD>10 or >5), the SNV burden increases to −44 and −127, respectively. Overall our results suggest that while the total burden of *de novo* mutations is stronger for STRs, the burden of inherited SNVs is stronger regardless of how conserved non-coding regions were defined.

## Discussion

Here, we presented SISTR2, a method for joint inference of the distribution of fitness effects (DFE) across a set of STRs. SISTR2 allows for improved inference of selection compared to SISTR at a broad range of repeat classes including those with low mutation rates or under only weak negative selection. We additionally leverage SISTR2 to refine estimates of STR mutation parameters, infer selection parameters across a diverse set of repeats, and estimate the relative burden of STR vs. SNV mutations in a typical genome.

We found that mutation and selection parameters are highly variable across STR classes. For example, we found that AT, AAG, AAAG, AAGG, and AGAT repeats have notably higher mutation rates than other STRs with the same repeat unit length. Further, while mutation rates for the majority of STRs scale linearly with repeat unit length, we found evidence that AAT repeats scale super-linearly with repeat length. Similarly, we estimate that STRs exhibit a wide range of selection coefficients, depending on the repeat unit and functional annotation of the repeat (**Fig. 3**).

We used SISTR2 to compare the expected overall fitness burden per individual of STR vs. SNV mutations. We considered both the burden of *de novo* mutations, as well as the burden due to inherited variants across the genome. The expected fitness of an individual STR mutation is estimated to be modest (10-fold less than a missense SNV). However, STRs are highly prevalent and experience a far greater number of mutations per individual. Thus, we estimate the *de novo* burden of STR mutations to be greater than that of SNVs. On the other hand, we found that the total burden of inherited variation is likely stronger for SNVs compared to STRs. We hypothesize that this may be because the total space of possible SNV mutations is much larger than that for STRs. Whereas only a small number of SNVs occur per generation, they accumulate over time as more sites are mutated. In contrast, STR mutations largely occur at a predetermined set of repeat elements already present in the genome. Further, STRs experience frequent “back” mutations which may reverse the effects of deleterious mutations in previous generations. Notably, our analysis was restricted to a subset of STR mutations with repeat unit lengths 2-4 bp that could be analyzed by SISTR2, and is thus likely an underestimate of the total burden of STR mutations. Still, our inferences of relative burden for both *de novo* and inherited variants are robust across a range of definitions of the sets of SNVs and STRs used for analysis.

Our study faced several limitations. First, one of the most significant challenges of our model is that we assume a single known optimal allele length at each locus and must analyze STRs with different optimal alleles separately. In practice, this optimum is unknown and is set to the modal allele. Further, it is not immediately clear why different STR loci with the same repeat unit would have different optimal lengths. Extending our model to relax this assumption or consider alternative models such as directional, rather than symmetric, mutation models is a topic of future study. Second, while our post-hoc analysis suggests inferred mutation and selection models fit the majority of STR classes well (**Supplementary Figs. 6-8, 10**), for some classes such as short trinucleotides or long AC and AAT repeats we could not identify a single best-fit model. This suggests for some repeat classes that our models do not completely capture properties of STR mutation, potentially biasing estimates of selection. Third, the burden of observed *de novo* STR mutations was found from analyzing Mendelian inconsistencies in trios using lymphoblastoid cell lines. These cell lines are known to accumulate mutations over time^16^, and thus mutation burdens computed from this dataset may be overestimated. To mitigate this risk, we computed the burden of STR mutations using several orthogonal strategies and across multiple trios (**Table 1**) and obtained similar results. Fourth, the model of selection within SISTR and SISTR2 assumes that mutations away from the modal allele are deleterious and that mutations act in an additive manner. Future work could examine whether more elaborate models, including positive selection and dominance effects might better fit the data. Finally, results here are based on allele frequencies from individuals of European descent. Computation of DFEs, and comparison of per-locus STR selection coefficients across populations, is an important topic of future work.

Overall, our findings suggest STRs are widespread targets of natural selection and that mutations at STRs contribute a substantial fitness burden in humans. Further, our results highlight important differences in mutation and selection across STR classes. These findings will inform future methods by enabling more accurate modeling of mutation and selection processes at STRs including improved inference of the impact of individual mutations that may contribute to evolution and disease risk.

## Methods

### SISTR2 mutation and selection models

The mutation, selection, and demographic history models are the same as those used previously in SISTR^7^. Briefly, the mutation model is described by four parameters: *µ*_*0*_ is the mutation rate of the optimal allele; *L* is the length-dependent mutation rate, such that the mutation rate of each allele x is determined as *µ*_*x*_ = *µ*_*0*_ *+ L*_*x*_; *β* is the length constraint, indicating the bias of long alleles to contract and short alleles to expand toward the optimal allele length; ρ is the step size parameter describing the geometric distribution from which mutation step sizes are drawn. The selection model assumes an optimal allele with fitness 1, with the fitness of other alleles decreasing with each number of units away from the optimal allele. We assume an additive fitness model, where the fitness of an individual is determined by summing the fitness of their two alleles at a locus. The demographic model is based on a published model of European population history^17^.

### Estimating the distribution of selection coefficients using SISTR2

SISTR2 uses ABC to estimate a distribution of *s* values (the distribution of fitness effects, or DFE) across a set of STRs (**Fig. 1a**). In this setting we assume the value of *s* at each STR is drawn from Γ(*a, b*) and learn posterior distributions for *a* and *b*. SISTR2 takes as input observed heterozygosities computed from allele frequencies for a set of STRs, plus a mutation model, selection model, demographic history model, and a prior distribution on the gamma distribution parameters *a* and *b*. The heterozygosity of locus *i* is defined as 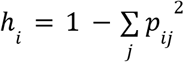 where *p*_*ij*_ gives the frequency of the *j*th allele at locus *i*. STRs for which a certain allele is fixed in the population have heterozygosity 0, whereas heterozygosity approaches 1 for highly variable STRs. To reduce computational complexity, SISTR2 first draws a random subset of 1000 loci from the input set of STRs (or use all the loci if there are less than 1000) and obtains the heterozygosity for each STR. It then repeats the following steps *z* (typically 50,000) times:

1. Draw gamma distribution parameters (*a, b*) from the priors, where *a* is drawn from a uniform distribution from 0 to 1 and *a* * *b* (mean value of the distribution) is drawn from a log normal distribution with a mean of 0.0003 and a standard deviation of 30.
2. For each STR *i* in the subset, draw *s*_*i*_ from the gamma distribution, simulate allele frequencies^5^ forward in time using this value of *s*_*i*_, and compute the heterozygosity of the resulting allele frequencies.
3. Compare the distribution of the simulated heterozygosity for all loci within the set to the distribution of the observed heterozygosity for all loci in the set by sorting both vectors, obtaining the difference vector, and taking the mean of the absolute value of the differences.

The *a,b* parameters that generate the top 1% of simulated heterozygosity distributions that are most similar to the observed distribution based on the mean of differences in step 3 above are accepted. We report the posterior estimate of the gamma distribution as the *a, b* pair with the median value of *a*\**b* (mean selection coefficient) out of all the accepted pairs. In practice to improve computational efficiency, we first generate a lookup table for each STR class (determined by its period and optimal allele) containing a list of selection coefficients and corresponding allele frequencies summary statistics. Then, for step 2 above, instead of simulating allele frequencies for each STR, we determine what class it belongs to and use the corresponding lookup table and randomly select a value of *s*_*i*_ sufficiently close to the *s*_*i*_ value drawn from the gamma distribution. For this, we rounded each *s* value to a single significant digit. Values <10^−5^ were rounded to 0. Two *s* values were considered sufficiently close if their rounded values matched.

### Validation of SISTR2 using simulated data

We validated SISTR2 using simulated datasets for six classes of STRs defined by their period (length in bp of the repeat unit) and optimal allele length. For each class we analyzed 12 pairs of gamma distribution parameters (*a, b*). For each class of STRs, for each gamma distribution parameter pair (*a, b*), we obtained the ground truth heterozygosity distribution obtained from simulating 1000 allele frequency distributions using the mutation model for that class and *s* values drawn from the gamma distribution characterized by *s* ~ Γ(*a, b*). Then, we performed ABC 20 times (each time with 2,000 simulations) to obtain 20 posterior estimates of the gamma distribution parameters as described above. Next, we calculated the mean *s* value (given by *a*\**b*) of each posterior estimate of a,b and plotted the mean of the 20 mean s values of estimated a,b parameters (**Fig. 1b**). To obtain the distribution of *s* values in different *s* bins for a given pair of parameters (*a,b*), we drew 1,000 *s* values using (*a,b*) and calculated the fraction of *s* values in each bin (**Fig. 1c**).

### Simulating genotyping errors

To evaluate the impact of STR genotyping errors on the results of SISTR2, we modified our simulation framework to add errors to simulated observed genotypes. We set the error probability for each observed allele to 0.1%. Each incorrect allele was set with 50% probability to be either one repeat larger or shorter than the true allele length. We then re-ran SISTR2 with these noisy observed genotypes as input (**Supplementary Fig. 3**).

### Genotyping STRs in European samples from the 1000 Genomes Project

Aligned whole genome sequencing CRAM files for the European (except Finnish) samples of the 1000 Genomes project were obtained from SRA accessions PRJEB31736 (unrelated samples) and PRJEB36890 (related samples). We excluded Finnish samples due to their unique population demographic history^9^.

GangSTR^8^ v2.4.5 was run on each sample separately with non default parameters --str-info str_info_file (see below), --bam-samps sample_id, --samp-sex sample_sex, and --grid-threshold 250. We generated an initial set of reference STRs for the hg38 assembly using Tandem Repeats Finder^18^ with the following parameters: match=2, mismatch=5, indel=17, maxperiod=20, pm=80, pi=10 and minscore=24. We then refined the STR reference set by applying a series of filtering steps. First, we removed repeats longer than 1Kb. Then, we kept a single repeat with the shortest motif length among those with identical start or stop coordinates. Compound and imperfect repeats were removed and any extra bases not matching the repeat motif were trimmed from both sides. Any duplicated repeats were discarded post-trimming. We then removed any repeats from the reference that did not have a minimum number of 10, 5, 4, and 3 copies for homopolymers, di-, tri- and tetra/penta/hexa-nucleotide repeats respectively. Finally, we filtered out any overlapping repeats if their motifs consisted of identical nucleotide types.

The file str_info_file contains the per-locus stutter parameters obtained by training the stutter model on 19 samples using a modified version of HipSTR v0.6.2 (https://github.com/mikmaksi/HipSTR) with non-default parameters --stutter-model-only (to skip genotyping), --chrom (to run separately for each chromosome), --min-reads 20, and --output-filters.

In the first merging step, mergeSTR^19^ v3.0.3 with non default parameter --vcftype gangstr was used to merge the VCF files of each sample into a unified VCF file for each sub-population (CEU, TSI, GBR, and IBS). The second merging step also uses mergeSTR v3.0.3 with non default parameter --vcftype gangstr to merge all of the sub-population level VCFs into one unified merged VCF for all European (except Finnish) samples. This merged VCF file was then filtered using dumpSTR^19^ v3.0.3 using non-default parameters --vcftype gangstr, --min-locus-callrate 0.8, --min-locus-hwep 0.00001, --filter-regions SEGDUP.bed, --filter-regions-names SEGDUP, --gangstr-filter-spanbound-only, --gangstr-filter-badCI, --gangstr-min-call-DP 20, --gangstr-max-call-DP 1000, --gangstr-require-support 2, and --gangstr-readlen 150. A list of segmental duplications (SEGDUP.bed) for hg38 reference genome build was obtained from UCSC table browser^20^. We then used statstr^19^ v3.0.3 with options --acount --numcalled to compute per-locus allele counts and record the number of genotyped samples per locus. We then filtered STRs with call rates <80%. We further excluded TRs with repeat lengths in hg38 <11 units for dinucleotides, <5 units for trinucleotides, and <7 repeats for tetranucleotides, since those repeats are typically not polymorphic. After filtering 86,327 STRs remained for analysis. The genomic annotation of each STR was assigned based on Ensembl^21^ build 92 for the GRCh38 reference genome.

### Computing weighted CDFs of selection coefficients

Cumulative distribution plots (**Fig. 3**) were computed by weighting results across DFEs inferred for all optimal allele lengths for each STR class. For each class, for each optimal allele we drew a number of *s* values from the learned DFE equal to the number of loci in that class. We then combined these randomly sampled *s* values across models for all optimal alleles, and computed cumulative distributions based on these combined values.

### Computing the burden of de novo STR mutations

We computed the reduction in fitness due to de novo STR mutations in two ways.

#### Predicted fitness burden

For each STR class *c* (motif/optimal allele length category), we used the model 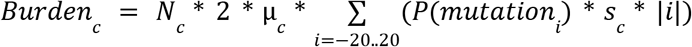 to compute the expected burden of *de novo* mutations for that class of STRs. *N*_*c*_ gives the number of loci in class *c*; *µ*_*c*_ gives the mutation rate of the optimal (modal) allele for class *c*; *i* represents all possible mutation lengths and ranges from −20 to 20 repeat units since mutations of larger sizes are extremely rare; *P*(*mutation*_*i*_) is the probability of a mutation resulting in a change *i* repeat copies, and is computed based on the step size parameter ρ (see SISTR^7^ for the full mutation model). *s*_*c*_ represents the mean selection coefficient for an STR in class *C* computeed by SISTR2 (**Supplementary Table 2**). The expected burdens from all the classes were then summed to obtain the genome-wide burden (**Supplementary Table 3**). We excluded STR classes with poor fit (KS Score<50, **Supplementary Table 2**) from downstream analyses.

#### Fitness burden based on directly observed *de novo* mutations

We additionally computed the observed burden of de novo mutations using three trios (children samples NA12864, NA10865, and NA10845 from the 1000 Genomes Project^9^). We first genotyped STRs using GangSTR as described above. We then used mergeSTR from the TRTools toolkit v4.0.0 to merge VCF files of individuals in each subpopulation. We applied call-level filtering on autosomal chromosomes of resulted VCF file using dumpSTR, from the TRTools toolkit v4.0.0 with parameters --min-call-DP 20, --max-call-DP 1000, --filter-spanbound-only, and --filter-badCI.

Then, filtered genotypes were subject to locus-level filtering with the parameters --filter-regions hg38_segdup_sorted.bed.gz, --filter-regions-names SEGDUP, --min-locus-callrate 0.8, and --drop-filtered to remove those loci that overlap with segmental duplications and those with low call rate. We then used statSTR from TRTools v4.0.0 to check consistency of parent genotypes with Hardy Weinberg Equilibrium (HWE) and used bcftools v1.12.14 to filter loci with a HWE p-value less than 10^−5^.

Finally, MonSTR^7^ v2.0 was used to call de novo mutations with non-default parameters --naive, --gangstr, --min-num-encl-child 3, --max-perc-encl-parent 0.05, --min-encl-match 0.9, --min-total-encl 10. We further removed de novo calls that were homozygous, removed loci that were biased toward expansion and deletion (two-sided binomial p-value < 0.05), removed mutations for which the *de novo* allele was supported by any reads in the parents and fewer than 5 reads in the child, and removed STRs for which more than 6 mutations across 568 total 1000 Genomes trios were observed implying that they are most likely error-prone STRs.

To compute the burden of each mutation, we scored the fitness of each allele as |*allele-optimal allele*|*\*s*, where *s* is the mean selection coefficient for that STR according to its class (motif/optimal allele category) based on SISTR2. Mutations at STR classes with poor model fit were excluded from analysis. The fitness effect of each mutation was computed as the difference in fitness between the new allele and the parent allele. Alleles resulting in fitness increases were excluded from results in **Table 1**.

### Computing the burden of inherited STR variants

The DFE for *de novo* mutations differs from the DFE for standing variation since the DFE for standing variation is biased toward more neutral values of *s*. For example, the *s* values for loci with inherited mutations that are a large number of repeat units away from the optimal alleles are likely not a random draw of *s* from the DFE inferred from SISTR2. Instead, these variants likely occur at loci that have values of *s* that are from the neutral portion of the DFE, otherwise those variants would have been removed by negative selection. Therefore, to compute the burden of inherited STR variants, we took a Bayesian approach that takes into account the frequency of a particular allele that is segregating in the population in the present-day.

For a given individual, for each STR variant at a locus with a SISTR2 score, we found which class *c* (motif/optimal allele category) the STR belongs to. Then, we calculated the reduction in fitness as |*allele* − *optimal allele*| * *s* To obtain the *s* for each inherited allele, 1,000 values of *s* from the gamma distribution corresponding to class *c* were drawn and allele frequencies were simulated using those values of *s*. For each of the 1,000 *s* values, if the simulated allele frequencies include the observed STR allele, the value of *s* and the STR allele’s frequency were recorded. For example, if we drew *s* = 0.001 and it gave the observed STR variant (15) and the allele 15 has a frequency of 0.2, we would record the pair (0.001, 0.2). From this, we get a table of (*s* value, frequency of desired STR allele) pairs. We then draw an *s* value from this table, with probability equal to the frequency of the mutant allele. For instance, if the table has (0.0001, 0.5), (0.001, 0.2), (0.01, 0.1), we would draw 0.0001 with a 5/8 chance, 0.001 with a 1/4 chance, and 0.01 with a 1/8 chance.

Burdens less than 10^−5^ were rounded down to 0 and greater than 1 were set to 1. Furthermore, if a variant was not found in any of the 1000 simulations, *s* was set to 0. Finally, for variants that were greater than 12 repeat units away from the optimal allele, s was automatically set to 0 since the simulations only contain 25 alleles in total. The fitness reductions for all STRs were summed to obtain the total STR burden for the individual. Mutations at STR classes with poor model fit (defined above) were excluded from analysis.

### Computing the burden of de novo single nucleotide mutations

To compute the burden of SNV mutations, we turned to previous estimates of the DFE for nonsynonymous mutations^13^ as well as for conserved non-coding mutations^14^. The mutational burden for each category was calculated as 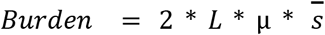, where *L* represents the mutational target size (i.e. number of sites that could be mutated) in a haploid genome, *µ* the per-base pair mutation rate, and 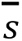, the mean selection coefficient, found from the inferred DFE (**Supplementary Table 4**). Mutational target sizes for noncoding regions were taken from Huber, *et al*.^*22*^. Given that the number of non-coding sites where mutations could have fitness effects is not precisely known, we explored a range of values ranging from 5% to 30% of non-coding sites being under selection (**Supplementary Table 4**).

### Computing the burden of inherited SNV variation

To compute the burden from standing genetic variation, we used an approach that links an estimate of the DFE to standing genetic variation via forward simulations. First, we used SLiM^23^ to simulate non-synonymous variation in a European population using a demographic history from Gravel et al 2011^24^ and a DFE from Kim et al 2017^13^. Similarly, we simulated non-coding variation using a DFE from Torgerson et al 2009^14^. From these simulations, we obtained the distribution of selective effects for heterozygous non-synonymous, homozygous non-synonymous, heterozygous conserved non-coding, and homozygous non-coding variants. Then, using data from the 1000 Genomes high coverage sequencing dataset^10^, we computed the number of heterozygous non-synonymous, homozygous non-synonymous, heterozygous conserved non-coding, and homozygous non-coding variants for each individual. In the real data, conserved non-coding mutations were annotated as having a CADD score >15 and annotations for non-synonymous mutations were obtained from files released by the 1000 Genomes project team (https://www.internationalgenome.org/data-portal/data-collection/30x-grch38).

For each mutation in each individual, we drew a selection coefficient from the simulated distribution of standing genetic variation, conditional on whether the mutation was conserved non-coding or non-synonymous and whether it was observed in a heterozygous or homozygous state. We computed additive fitness as 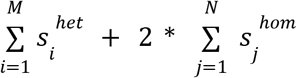 separately for both nonsynonymous and conserved non-coding, where *M* is the number of heterozygous sites in an individual and *N* is the number of homozygous sites in an individual.

## Data availability

Analysis is based on data from the 1000 Genomes Project available at SRA accession PRJEB31736.

## Code availability

SISTR2 source code and documentation can be found at: https://github.com/BonnieCSE/STRSelection.

## Acknowledgements

This study was supported by the Simons Foundation Autism Research Initiative (SFARI Grant #630705). M.G. was additionally supported in part by the Office Of The Director, National Institutes of Health under Award Number DP5OD024577 and NIH/NHGRI grant R01HG010149. K.E.L. was supported by the National Institutes of Health grant R35GM119856.

## Author contributions

B.H. designed and implemented SISTR2, designed and performed analyses of STR mutation and selection, and helped draft the manuscript. A.D. performed analyses of SNV mutation burden. N.M. and M.M. performed genome-wide genotyping of STRs and H. Z.-J. analyzed de novo STR mutations in the 1000 Genomes Project. K. E. L. and M.G. designed and supervised analyses and wrote the manuscript.

