## Supplementary Figures for "Genome-wide selection inference at short tandem repeats"

#### Supplementary Fig. 1

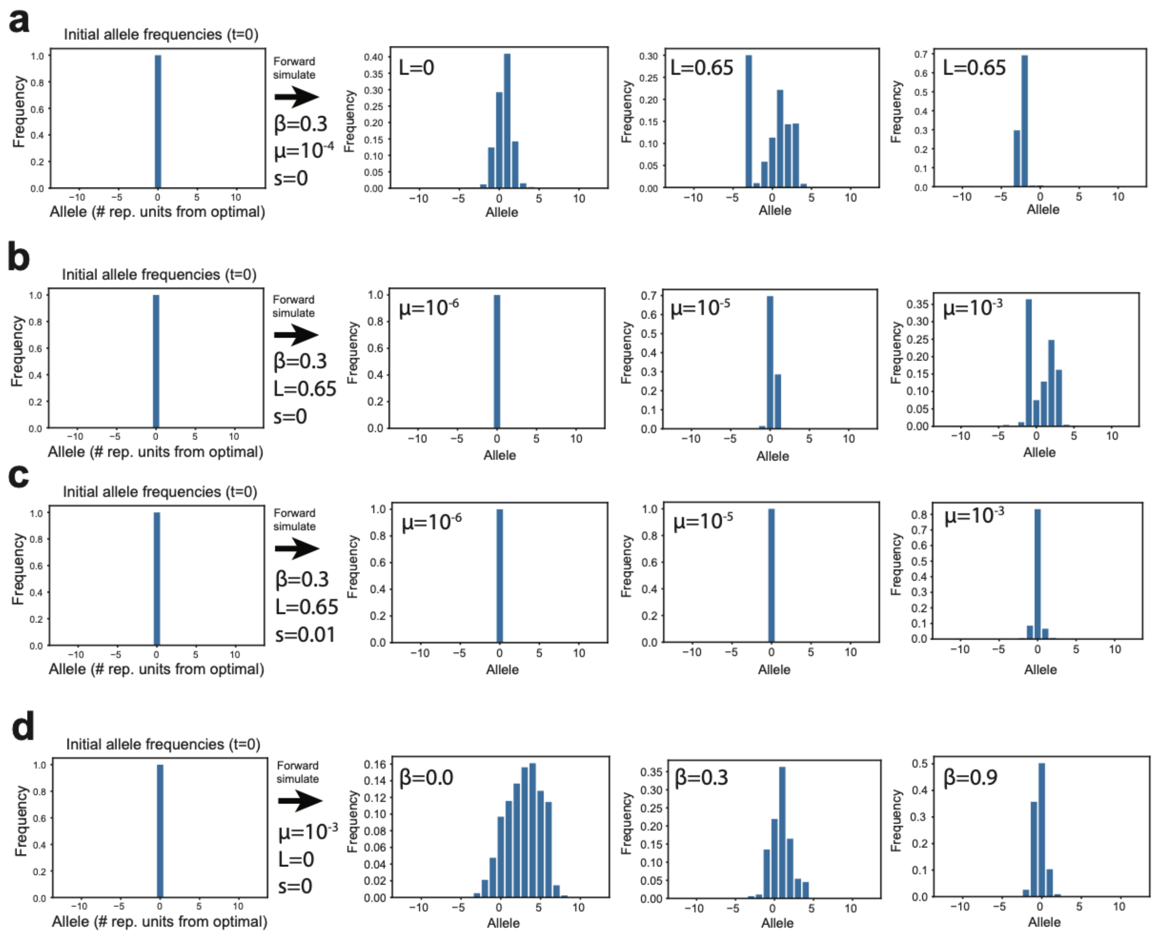


**Forward simulations demonstrate the effect of different parameters on allele frequencies. (a) Effect of *L* (length-dependent mutation rate parameter):** When *L* = 0, allele frequencies typically follow a bell-shaped curve, whereas when *L*>0, allele frequencies tend to follow a bimodal distribution. **(b-c) Effect of μ (per-generation mutation rate):** Given a value of *s*, as the mutation rate increases, the variability in the allele frequencies increases. When the mutation rate is low, it is difficult to distinguish between the effects of selection and low mutation at a single STR because there is little variability in the allele frequencies regardless of the selection coefficient. **(d) Effect of β (mutation size directional bias):** As **β** increases, so does the probability that an allele mutates toward the central allele. Thus, the resulting allele frequencies are more centered around the central allele and have lower variance. In each subplot, the x-axis gives allele size, in terms of the number of repeats away from the optimal allele (0). The y-axis gives the frequency of each allele in either the initial allele frequencies (left of the arrow) or forward simulated allele frequencies (right of the arrow). Results are based on representative replicates from running the forward simulation for *g*=20,000 generations.

#### Supplementary Fig. 2

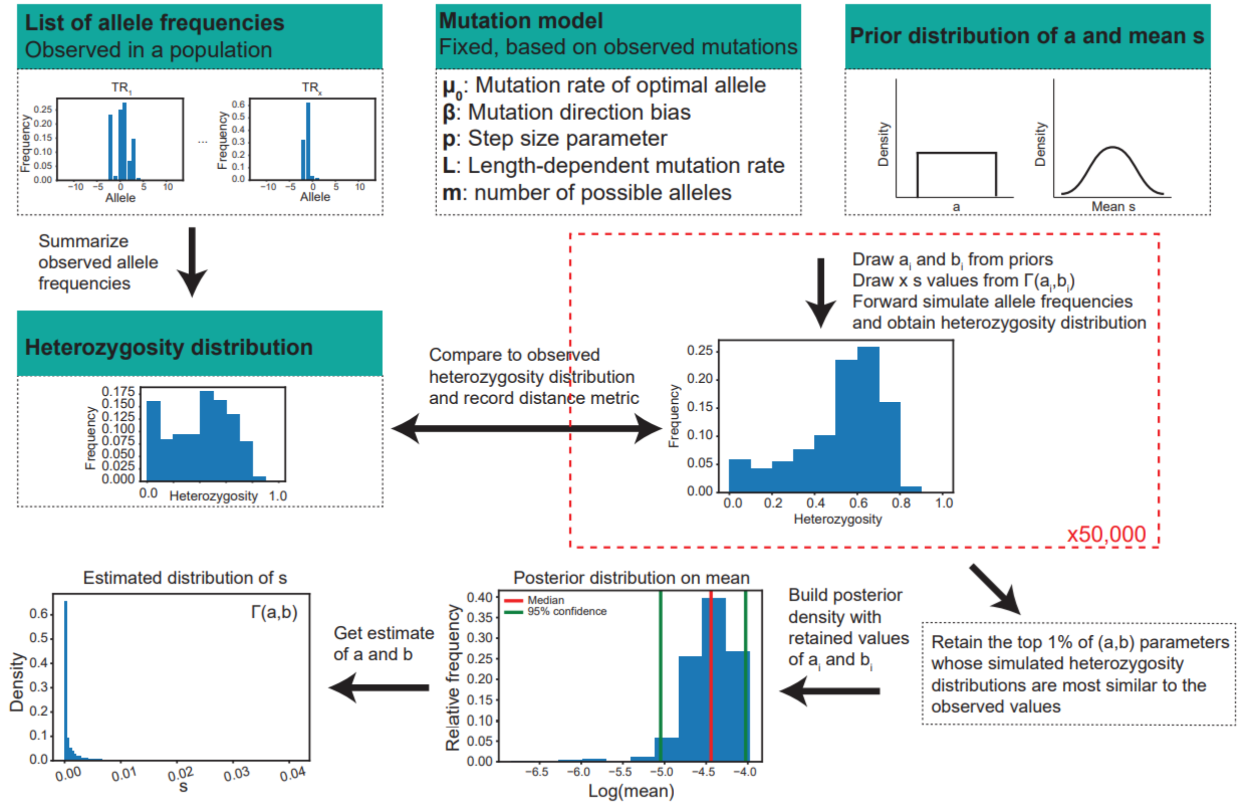


**Detailed schematic of SISTR2.** First, we start with a list of observed allele frequencies across a set of STRs and set the mutation model parameters based on the STR class the loci belong to (e.g. dinucleotide STRs with optimal allele length 11 repeats). Next, we perform approximate Bayesian computation (ABC) by drawing 50,000 (a,b) gamma distribution parameters from the prior, drawing x *s* values from each gamma distribution, simulating allele frequencies forward in time for each *s*, and comparing the resulting heterozygosity distribution with the empirical distribution using a mean of differences distance metric. The top 1% of (a,b) parameters with the most similar heterozygosity distributions to the observed values are accepted. Values of (a,b) from accepted simulations are used to construct the posterior distribution on the mean, which is calculated as s=a*b. The (a,b) pair with the median mean is used to obtain an estimated distribution of *s*.

### Supplementary Fig. 3

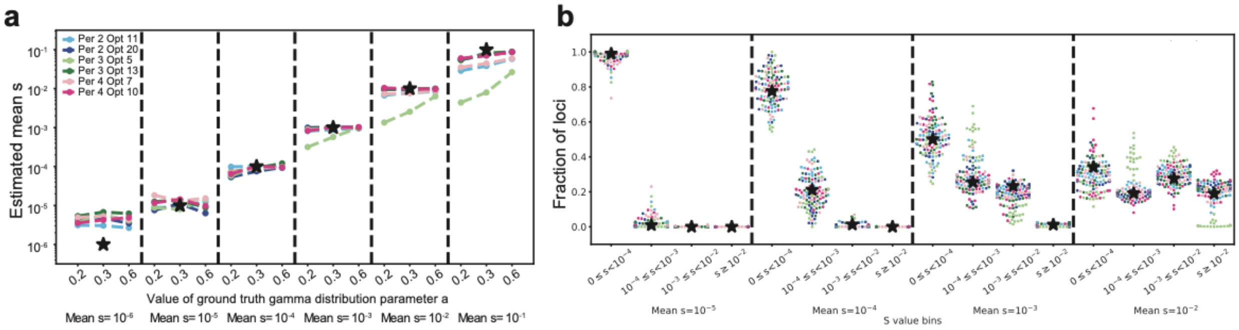


**Evaluating SISTR2 in the presence of genotyping errors. (a)** and **(b)** are the same as **Fig. 1b-c**, except “observed” allele frequencies contained simulated STR genotyping errors (**Methods**). Our results demonstrate that except in cases with extremely low underlying mutation rates (e.g. short trinucleotides), modest genotyping errors do not bias selection inferences.

#### Supplementary Fig. 4

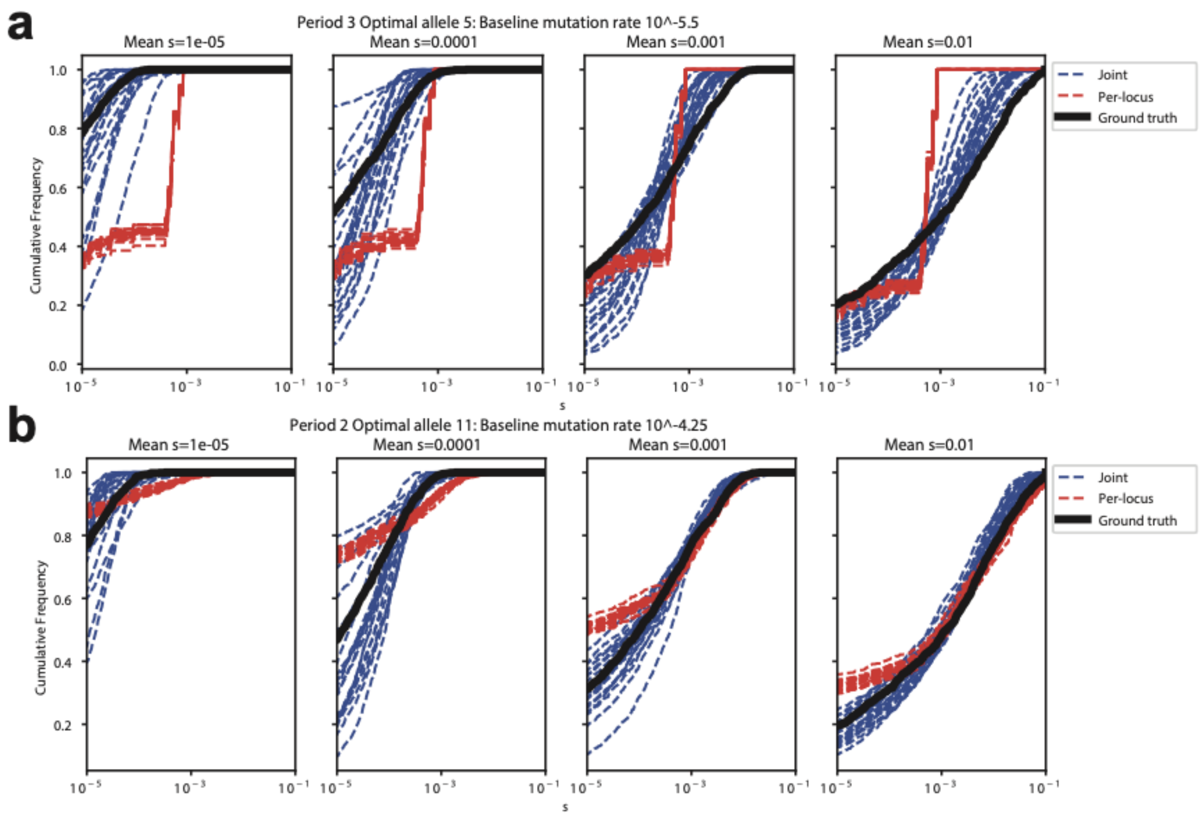


**Comparison of SISTR (per-locus selection inference) vs. SISTR2 (joint estimation across loci) on simulated data.** In each plot, the x-axis denotes the selection coefficient, and the y-axis shows the fraction of STRs in each category with selection coefficients greater than the x-axis value. **(a) Loci with a low baseline mutation rate.** Unlike the per-locus method (red lines), the joint method (blue lines) can estimate *s* for short loci with a low baseline mutation rate, such as trinucleotides with an optimal allele of 5 and a baseline mutation rate of $10^{-5.5}$. **(b) Loci with a high baseline mutation rate.** The per-locus method is better at inferring *s* for loci with higher mutation rates (e.g. dinucleotides with an optimal allele of 11 and baseline mutation rate of $10^{-4.25}$ as shown here) compared to loci with lower mutation rates. However, even at more highly mutable loci, the joint method is still more accurate than the per-locus method since it can distinguish between lower values of s. Overall, both the selection coefficient distributions obtained from the per-locus and joint methods are more concordant with the ground truth distribution as the mean *s* value increases.

#### Supplementary Fig. 5

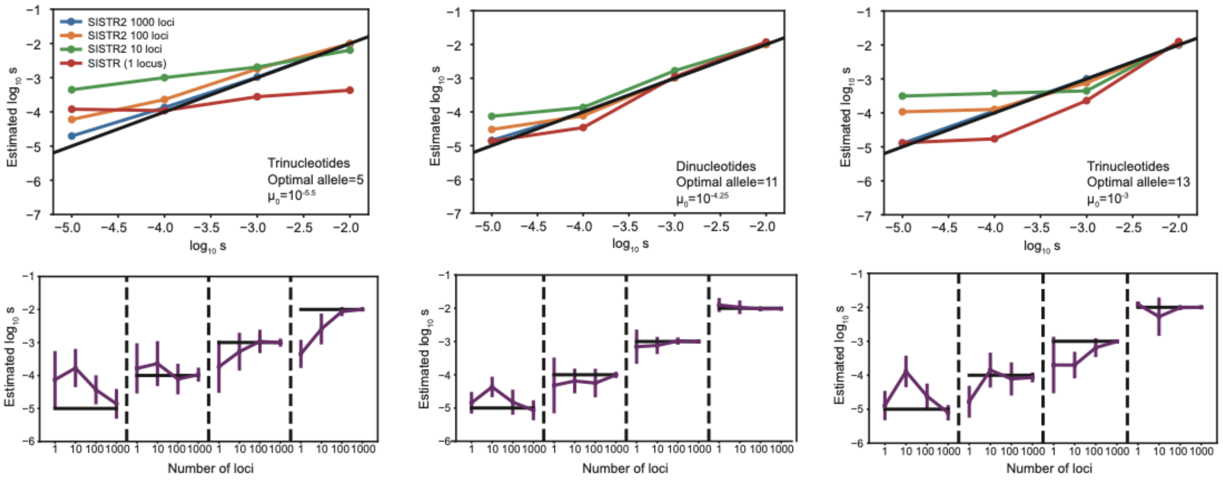


**Evaluation of SISTR2 selection inferences based on varying numbers of input STR loci.** We tested SISTR2 under various conditions (left=short trinucleotides with low mutation rates, middle=medium length dinucleotides with moderate mutation rates, right=long trinucleotides with high mutation rates). For each condition, we evaluated SISTR2's selection inferences as a function of the number of STR loci used as input. In the top panels, the x-axis gives the simulated mean value of *s* and the y-axis gives the inferred mean value across 20 estimates. Colors denote the number of loci used for inference (red=1, green=10, orange=100, blue=1000). Bottom plots show the mean +/- 1 s.d (purple) across the 20 simulations in each condition. True mean values of *s* are given by black horizontal lines. For all simulations shown, gamma distribution parameter *a* (i.e. the shape parameter) was set to 0.6.

#### Supplementary Fig. 6

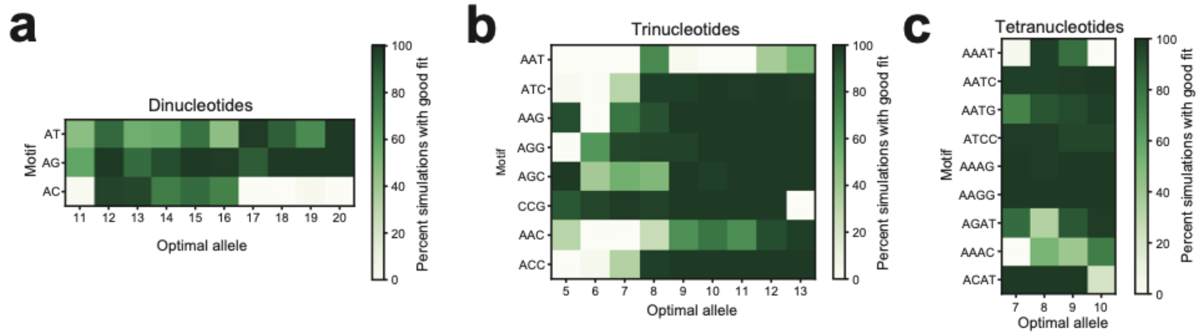


**Assessing the goodness of fit of maximum likelihood estimates of mutation models.** For each STR class, corresponding to a particular repeat unit (x-axis) and optimal allele (y-axis), we determined the mutation model that maximizes the likelihood (**Fig. 2**) and assessed the goodness of fit with observed data. For each class, we simulated 100 datasets based on the inferred mutation model, and compared the distribution of heterozygosities for simulated loci to those from observed loci using a KS test. Each heatmap cell shows the percent of simulation rounds with KS test p-value>0.05, indicating the two distributions are similar and the model fits well.

#### Supplementary Fig. 7

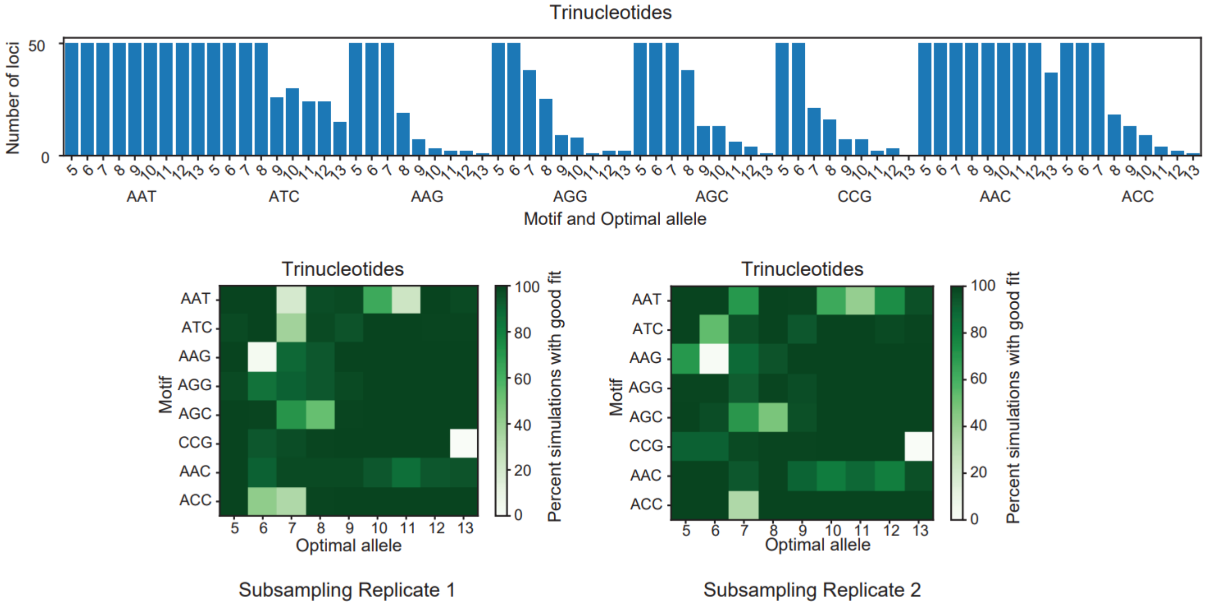


**Assessing mutation model goodness of fit on subsampled data.** To reduce differences in power across different STR classes to detect differences between observed vs. simulated heterozygosity distributions, we repeated the goodness of fit analysis (**Supplementary Fig. 6**) but including at most 50 STRs sampled from each class. Top panels show the number of STRs included in each class. Bottom panels are the same as in **Supplementary Fig. 6**).

#### Supplementary Fig. 8

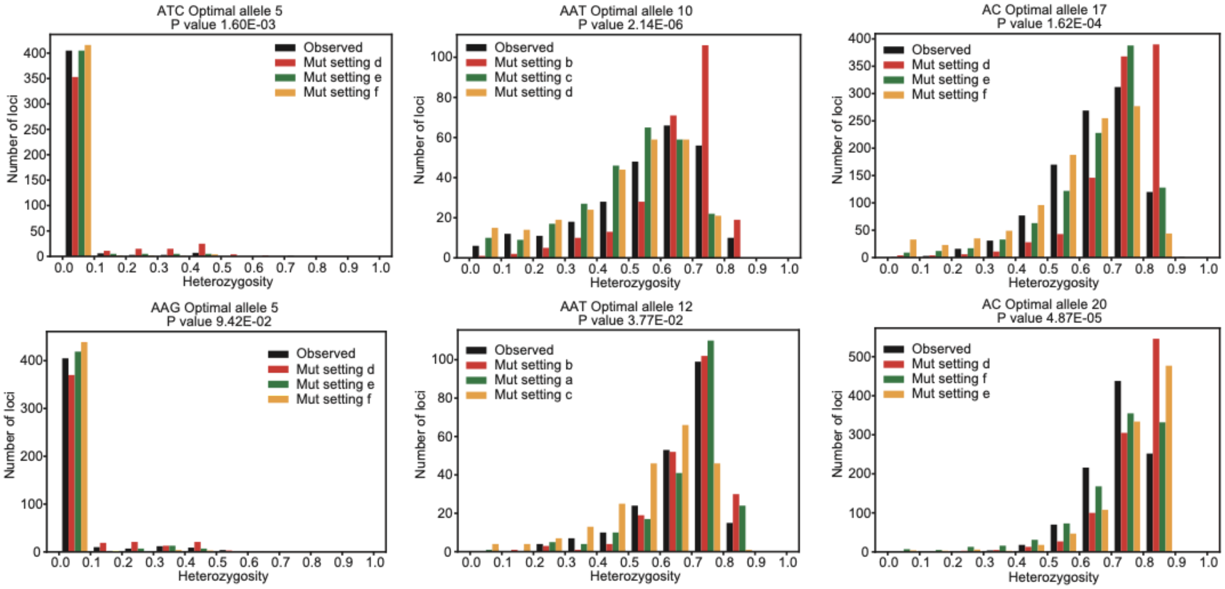


**Visualization of observed vs. simulated heterozygosity distributions.** For classes of loci with a low percentage of simulations that fit the maximum likelihood mutation model well, we plotted the observed vs. simulated heterozygosity distributions for a range of models to visually assess their similarity. The observed heterozygosity distribution is in black, the maximum likelihood mutation model is in green, and other mutation models for comparison are in red/orange. Visually, the maximum likelihood mutation model results in simulated heterozygosity distributions close to observed distributions, even for cases where the KS test indicated a poor fit.

#### Supplementary Fig. 9

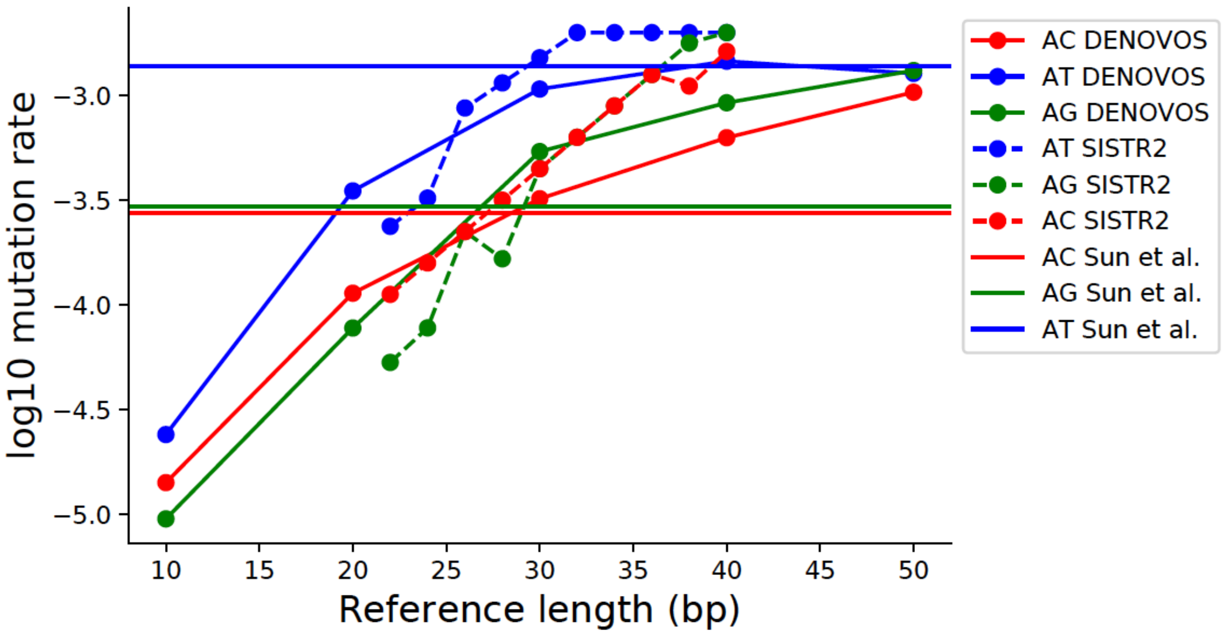


**Comparison of repeat unit-specific dinucleotide mutation rates inferred from SISTR2 versus de novo mutations.** We compared the dinucleotide mutation rates inferred by SISTR2 (dashed lines) to those inferred from *de novo* mutations based on WGS of the Simons Simplex Collection (SCC) dataset[1] (solid lines with points) and from *de novo* mutations based on capillary electrophoresis[2] from Icelandic individuals (solid horizontal lines). The SISTR2 inferred mutation rate for each motif/optimal allele combination was set as the median mutation rate in the posterior of accepted mutation models from the feasible mutation parameters analysis (**Fig. 2**). In all datasets, AT repeats have higher mutation rates than AC or AG repeats. Additionally, inferences from SISTR2 and in SSC show AC repeats mutate faster than AG repeats for shorter but not longer repeat tracts.

### Supplementary Fig. 10

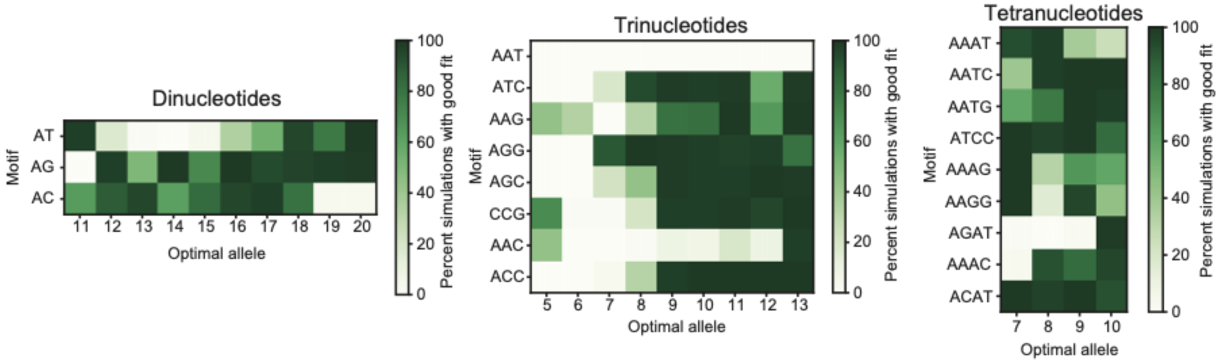


**Goodness of fit assessments of the SISTR2 motif analysis scores.** For each class of loci (i.e. optimal allele/repeat unit pair), 100 heterozygosity distributions were simulated using the distribution of selection coefficients inferred by SISTR2. Then, a KS test was used to compare the simulated distributions to the observed distributions. Each heatmap cell shows the percent of simulation rounds with KS test p-value>0.05, indicating the two distributions are similar and the model fits well. These values are reported as the KS Score in **Supplementary Table 2**.

#### Supplementary Fig. 11

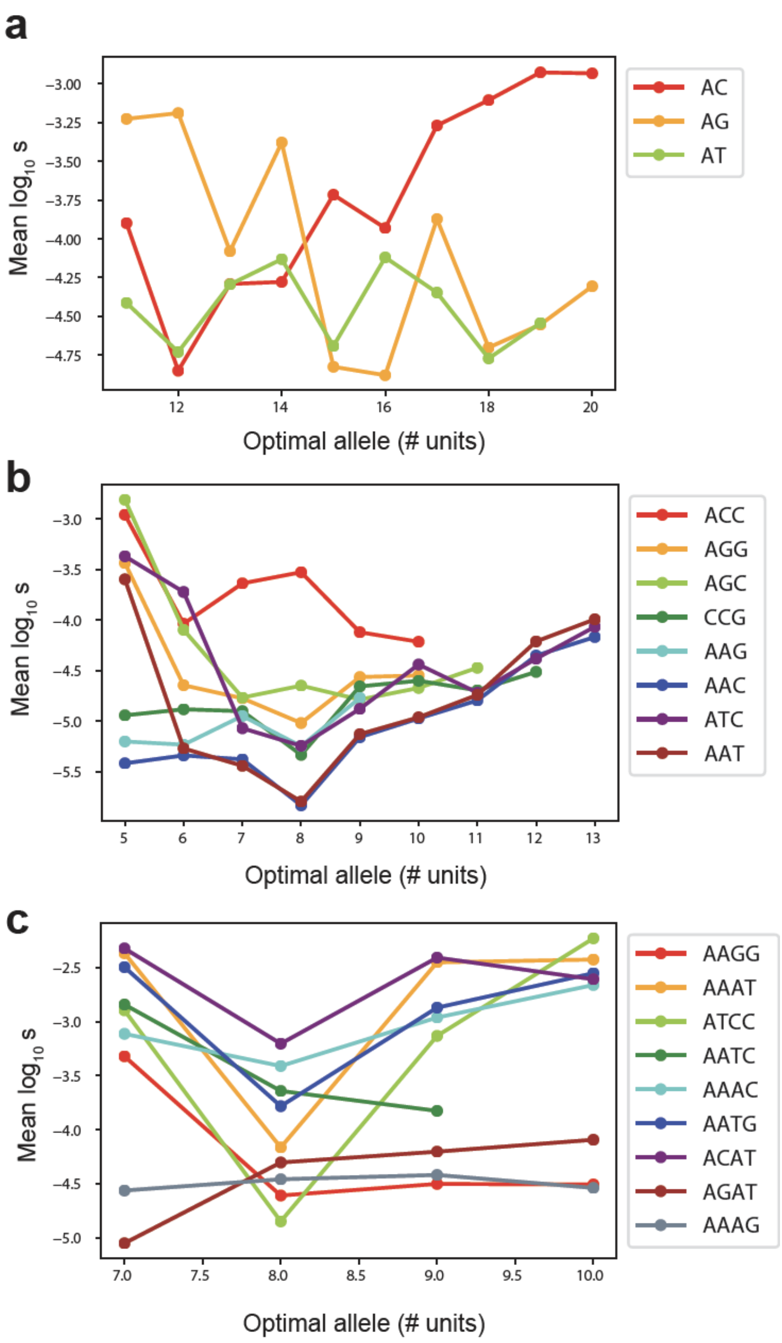


**Mean selection coefficient for each repeat unit and optimal allele length.** In each panel, the x-axis gives the optimal allele length and the y-axis gives the median inferred selection coefficient. Each color denotes a different repeat unit sequence. Categories computed based on fewer than 10 loci were excluded from this analysis.

#### Supplementary Fig. 12

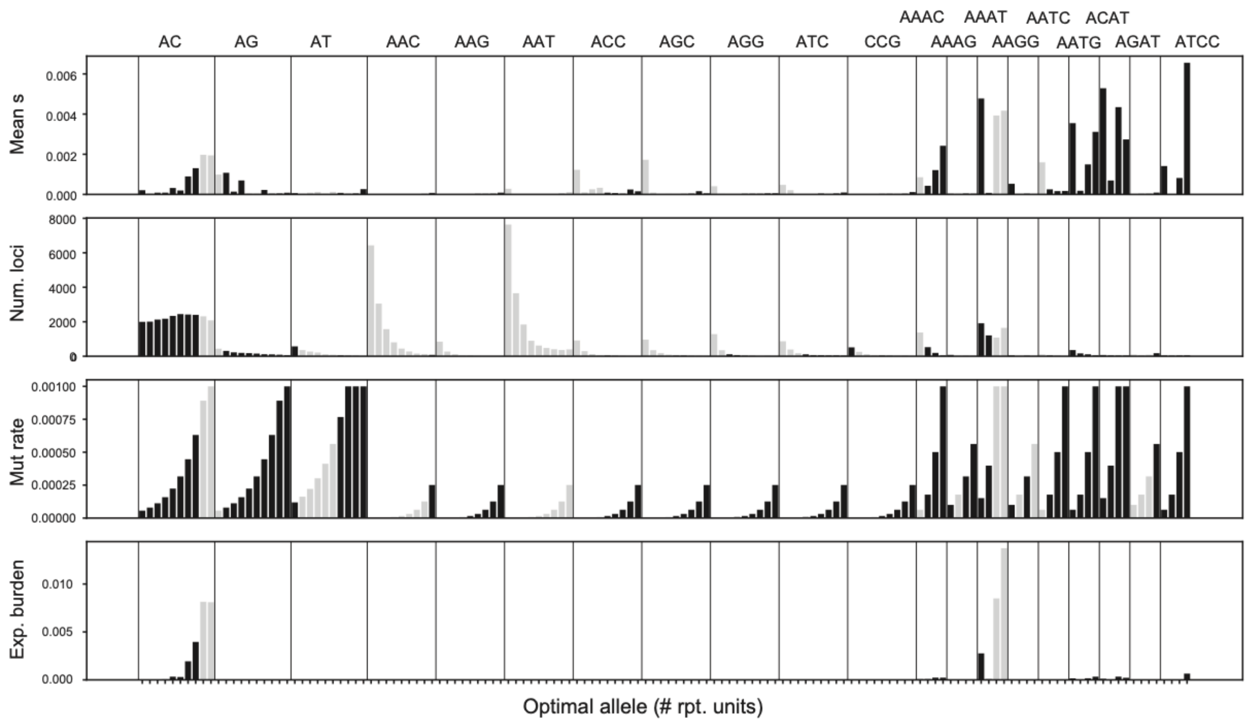


**Mutation/selection parameters and expected burden of mutations for each STR class for *de novo* mutations.** The x-axis shows different STR classes based on the repeat unit (annotated at the top) and optimal allele length. Optimal allele lengths range from 11-20 repeats (dinucleotides), 5-13 repeats (trinucleotides), and 7-10 repeats (tetranucleotides). Top panel: mean selection coefficient per mutation in each class. Second from top: number of STR loci considered in each class. Second from bottom: mean mutation rate for each class. Bottom: expected burden of *de novo* mutations for each class. STR classes excluded from burden calculations due to low fit are colored in gray.

#### Supplementary Fig. 13

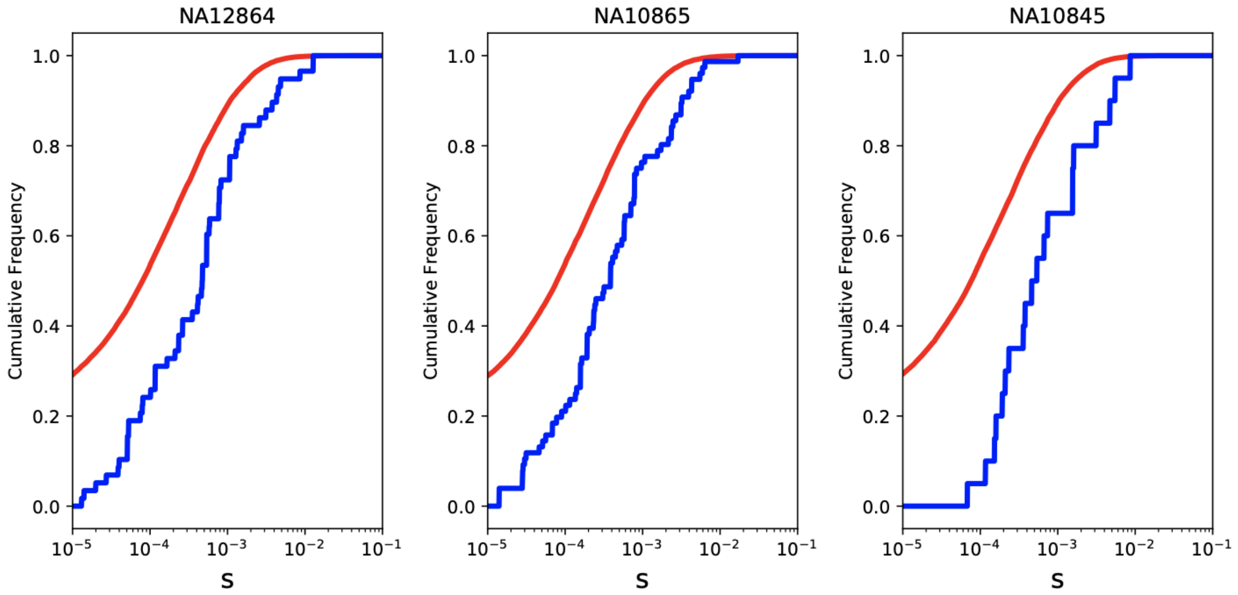


**Burden of de novo vs. standing variation in three children.** For each sample, we determined the selection coefficient for each non-optimal inherited allele (red lines) (detailed procedure described in **Methods**) and each allele resulting from a *de novo* mutation (blue lines) using SISTR2. *De novo* mutations resulting in optimal alleles were excluded. Lines show the cumulative frequencies of selection coefficients for each category.
